# Spatiotemporal 3D image registration for mesoscale studies of brain development

**DOI:** 10.1101/2021.05.12.443839

**Authors:** Sergey Shuvaev, Alexander Lazutkin, Roman Kiryanov, Konstantin Anokhin, Grigori Enikolopov, Alexei A. Koulakov

## Abstract

Comparison of brain samples representing different developmental stages often necessitates registering the samples to common coordinates. Although the available software tools are successful in registering 3D images of adult brains, registration of perinatal brains remains challenging due to rapid growth-dependent morphological changes and variations in developmental pace between animals. To address these challenges, we propose a multi-step algorithm for the registration of perinatal brains. First, we optimized image preprocessing to increase the algorithm’s sensitivity to mismatches in registered images. Second, we developed an attention-gated simulated annealing (Monte Carlo) procedure capable of focusing on the differences between perinatal brains. Third, we applied classical multidimensional scaling (CMDS) to align (“synchronize”) brain samples in time, accounting for individual development paces. We tested this multi-step algorithm on 28 samples of whole-mounted perinatal mouse brains (P0 – P9) and observed accurate registration results. Our computational pipeline offers a runtime of several minutes per brain on a personal computer and automates brain registration tasks including mapping brain data to atlases, comparison of averaged experimental groups, and monitoring brain development dynamics.

## INTRODUCTION

The development of algorithms for transforming similar images into common coordinates (*registration* algorithms) was initially driven by the need to register low-resolution medical images, such as fMRI data (Oliveira and Tavares, 2014). Since then, substantial progress has been made leading to the development of optimized image registration software that is now used in clinical practice. The emergence of whole-brain staining and imaging methods has yielded whole-brain datasets with a single-cell resolution, which require high levels of registration precision (reviewed by Susaki and Ueda, 2016). Large datasets containing whole-brain imaging data collected by 3D microscopy labs require high-throughput and precise automatic registration to brain atlases. In this work, we build upon two key existing image registration approaches to produce a set of image registration algorithms that could enable mesoscale-level studies of brain development.

The two main approaches to image registration are *feature-based* and *free-form* registration. *Feature-based registration* allows users to specify pairs of landmarks in two images, which are used in alignment (Lerios et al., 1995). With increasing number of features, this procedure can reach user-defined levels of precision. Overall, feature-based registration is a flexible and computationally tractable approach (Oliveira and Tavares, 2014), however, it may require significant input from users. To increase the throughput of feature-based registration, multiple research groups have worked towards automatic feature extraction methods. One successful approach involves searching for features in the frequency domain. Fourier transformations (Hughes, 1992) and wavelets (He et al., 1994) were used to identify independent image components, which were then registered separately. Feature-based registration techniques have proven to be efficient on low-resolution medical images (Andreetto et al., 2004).

*Free-form methods* use a different, complementary concept of image registration (Sederberg and Parry, 1986). Instead of aligning hand-picked features, free-form registration methods optimize spatial transformation to maximize the similarity between two images. Following Sederberg and Parry (1986), one may conceptualize free-form registration as deforming a brain sample together with the transparent agarose cube in which it is embedded to increase similarity integrated over the entire image. The similarity between samples is usually evaluated using measures such as correlation, mutual information, or L2 norm of differences; as a result, the outcomes of registration depend on data preprocessing (Klein et al., 2009a). In successful applications, alignment is typically preceded by image smoothing (Niedworok et al., 2016) and involves converting raster images to vector fields (Allen Brain Institute, 2017).

Free-form registration methods gained popularity due to their ease of automation. Residual discrepancies between samples, inevitably arising from the methods’ insensitivity to small displacements (Viergever et al., 2016) are shown to not affect major brain regions (Niedworok et al., 2016). These recent free-form registration algorithms are largely efficient and are widely used for mapping high-resolution brain data to brain atlases.

In this work, we sought to combine the automation of freeform approaches with the precision of feature-based methods to align multiple brains across different developmental stages and to reconstruct mesoscale developmental dynamics of the perinatal mouse brain. We have determined what image features are critical for proper and efficient brain registration and used those in a free-form transformation routine. We introduced routines for estimating the real developmental age for individual samples and for detecting changes occurring in the samples over time (*Figure 1*). We applied our procedures to 28 samples of perinatal mouse brains dated P0 – P9. We show that our algorithms address challenges specific to the registration of perinatal mouse brains.

**Figure 1.**
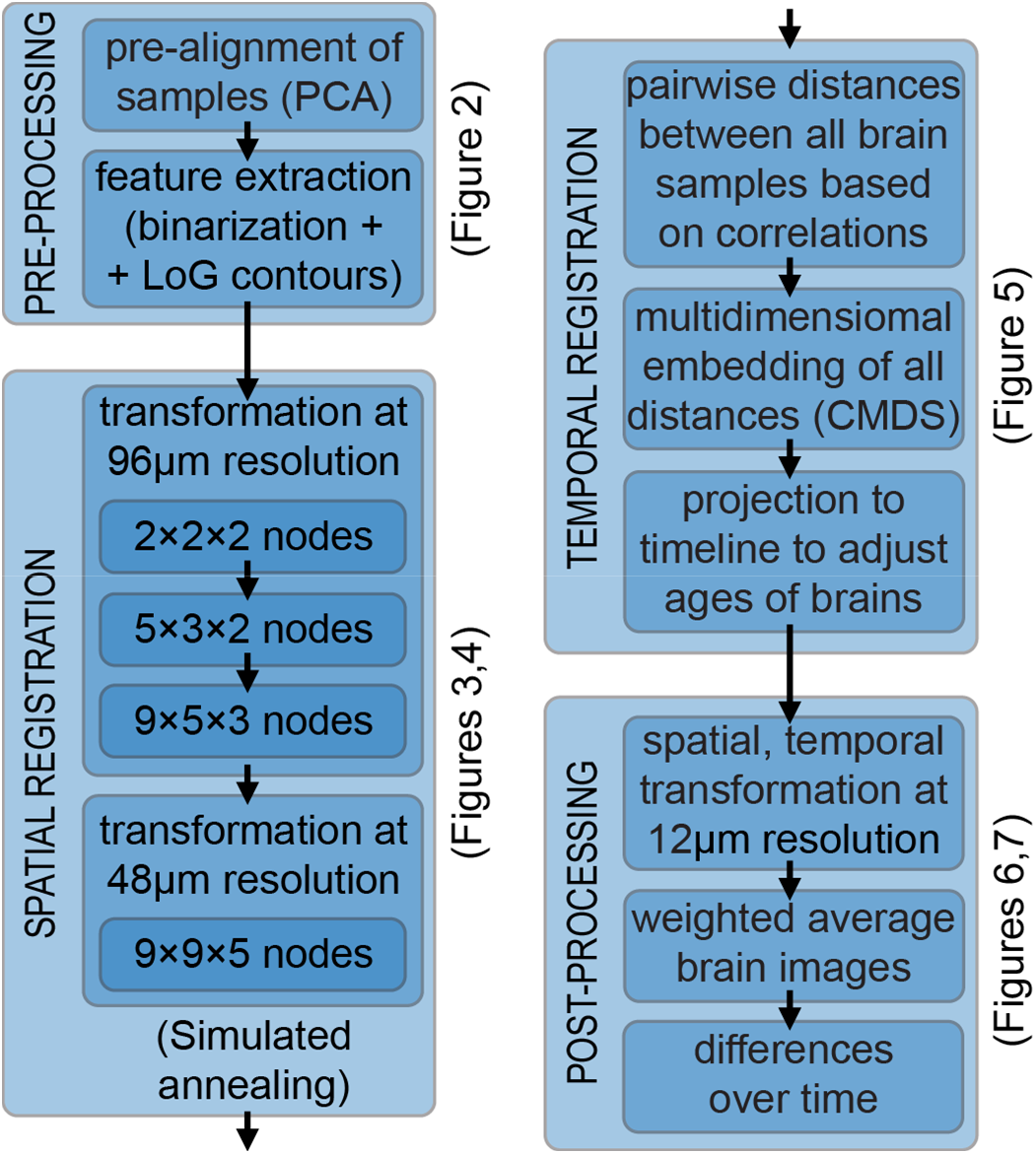
Flow chart of our algorithm for uncovering mesoscale developmental dynamics in the brain. To register brain samples, we start with preprocessing which includes pre-alignment using the principal component analysis (PCA) and extracting the image features to be registered. These features include brain region contours (extracted with Laplacian of Gaussian (LoG) filter) and the overall brain shapes (represented using the binary mask). We then perform spatial registration of brain samples using piece-wise linear deformations and an attention-gated simulated annealing Monte Carlo algorithm. We further register brain samples in time by building a multidimensional embedding where distances between brains are defined by their pairwise dissimilarities (classical multidimensional scaling; CMDS). We project the axis of maximum variance on the timeline, thus obtaining the adjusted ages for each brain sample. We finally display the results by combining the registered brain images with the weights reflecting their adjusted ages. We also compute differential images reflecting short-term developmental dynamics and color-code identified differences.

## METHODS

### Data import

We downsampled the 3D images of the brains to a uniform resolution of 12×12×12 μm per voxel. We applied histogram equalization within the XY-slices of these images to equalize the signal intensity along the Z-axis in the samples. To remove the background signal, we selected a threshold to best separate the signal inside and outside the sample (~0.01 of the maximum signal). We suppressed the signal below this threshold. To pre-align the 3D images, we used the principal component analysis (PCA) (Wold et al., 1987) (*Figure 2A*) on binarized versions of the images (1 for voxels >0.01 of maximum intensity, *Figure 2B*). We chose an orientation of a sample along the PCA axes manually based on correlations between binarized samples.

**Figure 2.**
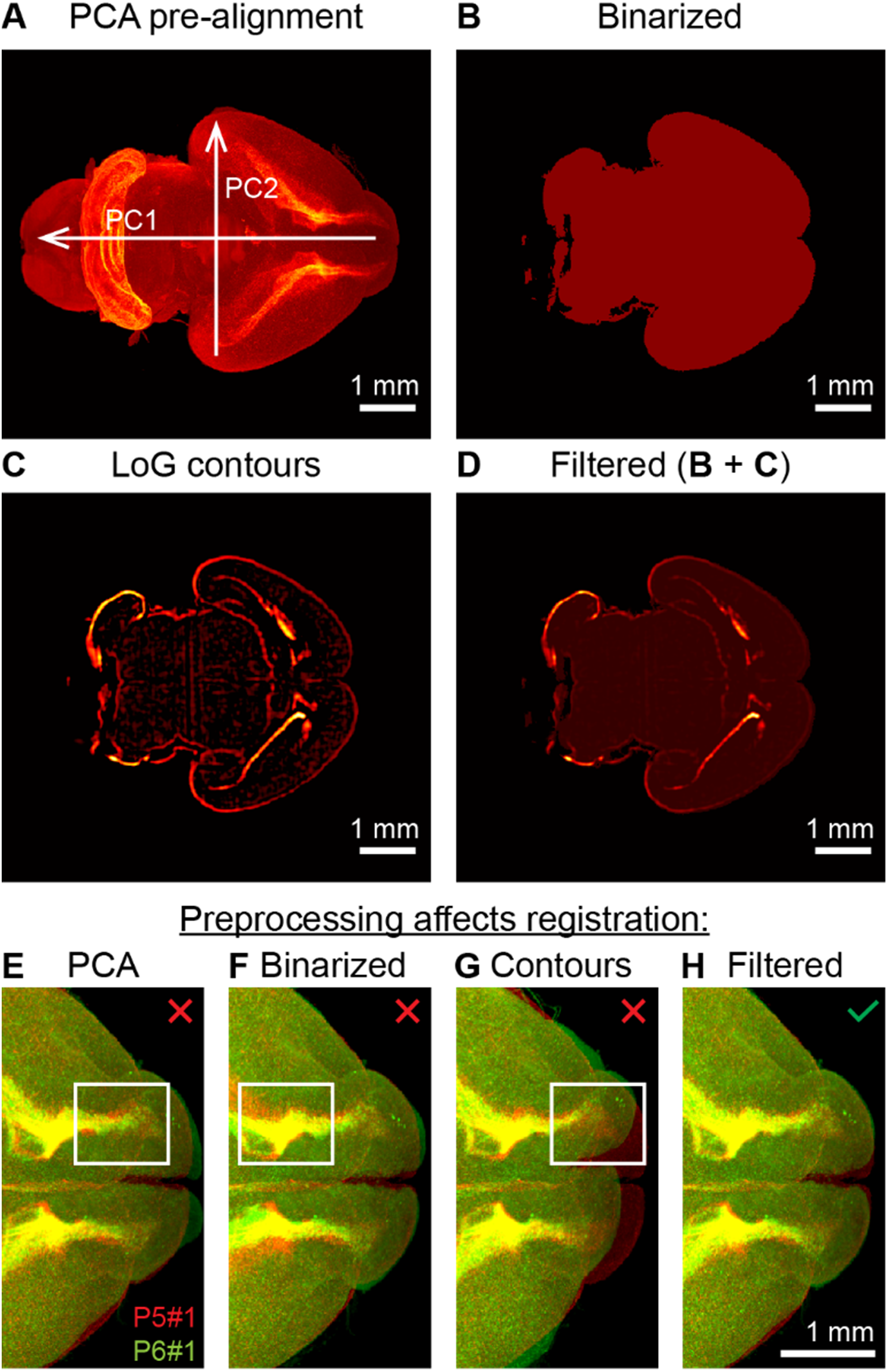
3D image preprocessing steps improving brain registration. (A) Brain sample pre-alignment via rotation and scaling using Principal Component Analysis (PCA). (B) Cross-section of the binary mask of the brain sample. (C) Contours of the brain regions extracted with Laplacian of Gaussian (LoG) filter. (D) Filtered image; a weighted sum of (B) and (C). (E-H) Registration of two brains taken at P5 and P6 using individual image preprocessing methods as indicated. Registration of raw images (E) results in slight mismatches between Rostral Migratory Streams (RMSs). Registration of binary masks (F) results in a mismatch of the inner structure. Registration of brain region contours (G) may not converge and results in large mismatches between samples. The combination of binary mask and contours (H) solves the problems of (E-G) yielding a higher quality registration.

### Feature extraction

We extracted contours of brain regions (*Figure 2C*) using the Laplacian of Gaussian (LoG) filter (Huertas and Medioni, 1986). In the LoG filter, we set the blur radius large enough to average out individual cells in the samples, yet small enough to preserve shapes of brain regions. We found that the blur radius of 5 voxels provided a good compromise for our images. To accelerate computation, we performed the LoG filtering in frequency domain. We normalized the intensity of voxels in the contour images (produced with the LoG filter) to have an average value of 1 over non-zero voxels (*Figure 2C*). We added the normalized contour images to the binarized images (*Figure 2B*) to obtain the *filtered* images (*Figure 2D*).

### Spatial registration

For each 3D image of a brain sample, we defined a set of spatial transformations ranging from coarse to fine. Each transformation was defined by displacements of nodes in a regular grid spanning the entire 3D image; grid spacing specified the coarseness of transformation (*Figure 3A*). Displacements of individual voxels in the 3D image were computed via linear interpolation of the grid node displacements (*Figure 3B*). To transfer signal intensity from original to new coordinates we used linear interpolation (Bourke, 1999) of signal intensities. We optimized the node displacements to maximize the objective function defined as the Pearson correlation of filtered (*Figure 2D*) brain images minus 1/1000 of deformation energy. To estimate the deformation energy, we triangulated the transformed 3D image using the grid nodes, then computed the L1 norm of the volume changes in the triangulation compartments.

**Figure 3.**
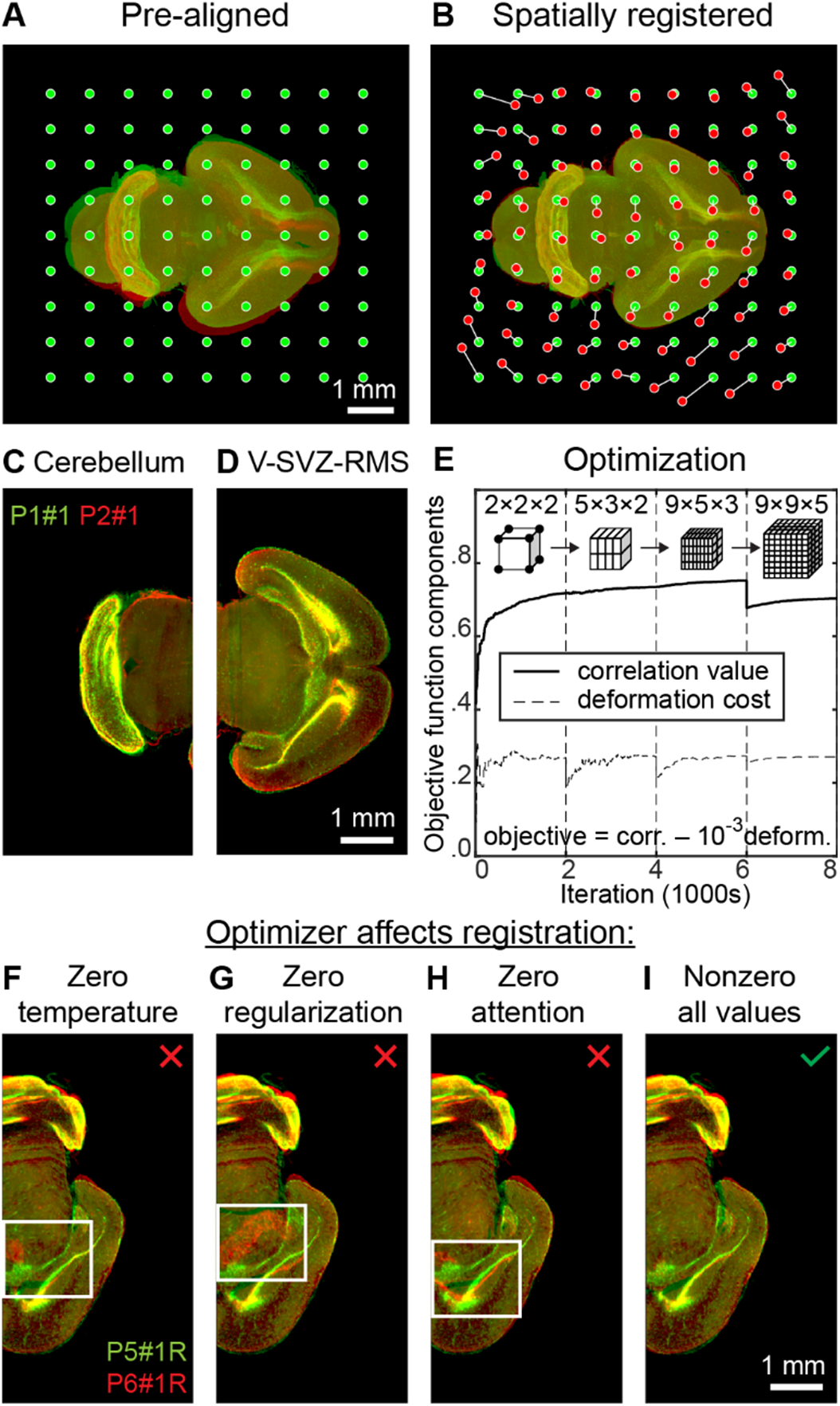
The simulated annealing algorithm allows registering dissimilar developing brains. (A) Two brain samples before alignment and the nodes of transformation grid. (B) Registered brain samples and displacements of the nodes of the transformation grid. (C) Registered cerebellum and (D) ventricular-subventricular zone (V-SVZ) and rostral migratory stream (RMS). (E) Pearson correlation (similarity) between two filtered brain images (solid line) and deformation cost for the registered brain (dashed line) throughout the simulated annealing registration. The objective function maximized with simulated annealing is a weighted difference between correlation value and deformation cost (1:1000). Simulated annealing is performed in 4 stages, separated by vertical dashed lines, corresponding to different numbers of cells in the transformation grid (2×2×2; 5×3×2; 9×5×3; 9×9×5) and different image resolutions, defined by linear voxel size (96μm; 96μm; 96μm; 48μm). The drop of similarity at iteration 6000 is due to refinement of the voxel size (96μm → 48μm). (F-I) Registration of P5 and P6 brains with different optimizers. (F) Registration using a greedy algorithm may result in mismatches of some brain structures. (G) Registration with unregularized simulated annealing may diverge, resulting in large-scale discrepancies between source and target 3D images. (H) Simulated annealing without attention gating, i.e. choosing nodes with equal probability, converges slower than (I) simulated annealing with attention gating, i.e. prioritizing nodes with poorly aligned surroundings.

### Optimization

To optimize the 3D image transformation, we attempted to displace nodes of the transformation grid one by one using the Simulated Annealing (SA) algorithm (Van Laarhoven and Aarts, 1987) (*Figure 3E*). We sampled the magnitude of each attempted displacement from a zero-mean Gaussian distribution. The probability for a displacement to be accepted was defined by the equation *p* = min(exp(Δ*E* / *t^o^*),1) which depended on a displacement-related change in the cost function, Δ*E*, and on *temperature t^o^* (Van Laarhoven and Aarts, 1987).

### Optimization procedure

To accelerate the convergence of the algorithm, we were selecting the nodes to be displaced with probability proportional to the L1 difference between the source and target images, both filtered, within the cells of the transformation grid adjacent to the node. The coarseness of alignment was defined by the spacing of the transformation grid. The image resolution and the number of nodes in the transformation grid were refined in four stages of alignment, ranging from coarse to fine (Table 1; *Figure 3E*). We updated the grid spacing at the beginning of each stage of alignment by adding new nodes to the transformation grid in accordance with Table 1. We defined the displacements for newly added grid nodes using linear interpolation of the existing displacements and set the magnitude of the average attempted displacement equal to 20% of the new grid spacing. At the beginning of each stage of alignment, we also initialized temperatures to be used in determining acceptance of displacements (Table 1). We were updating these parameters throughout each stage of alignment. Temperatures t^o^ were decreased exponentially; their values at the end of a stage equaled to 1/30 of their values at the beginning of the same stage. The magnitude of the average attempted displacement evolved depending on the success of previous displacement attempts. Specifically, we divided/multiplied the magnitude of the average attempted displacement by 0.99 at each successful/unsuccessful SA iteration.

**Table 1.** Stages of spatial alignment

| Stage | I coarse | II | III | IV fine |
| --- | --- | --- | --- | --- |
| Iterations | 2000 | 2000 | 2000 | 2000 |
| # of nodes | 2x2x2 | 5x3x2 | 9x5x3 | 9x9x5 |
| Resolution | 96 μm | 96 μm | 96 μm | 48 μm |
| Initial $t^0$ | $10^{-3}$ | $10^{-3}$ | $10^{-4}$ | $10^{-5}$ |
| Final $t^0$ | $10^{-3} / 30$ | $10^{-3} / 30$ | $10^{-4} / 30$ | $10^{-5} / 30$ |

Each stage of alignment was also characterized by the resolution to which we downsampled the 3D brain images on that stage. We defined resolution by the linear size of a voxel, the same for all 3 dimensions (Table 1). Upon completion of all stages of alignment, we applied final transformations, obtained on downsampled images and defined by displacements of the grid nodes, to full resolution images (*Figure 3C, D*).

### Temporal registration

To provide additional validation of our approach, we applied our procedures independently to different hemispheres of the same brain. To this end, we split each 3D image of a brain into two separate images of hemispheres, which doubled the number of samples in our dataset. To split an image of a brain close to symmetrically, we used the following procedure. First, we aligned each brain to its mirror image within the volume. This yielded a transformation T. Second, we transformed the original brain using ½ of this transformation, i.e. T/2. This procedure placed the plane of the brain’s symmetry to the middle plane of the 3D volume. Finally, we split the brain into two hemispheres using the middle of the 3D volume as a separatrix. This procedure yielded two images of hemispheres per brain sample.

Then we aligned the right hemispheres of different brains to each other. To set the order of this alignment, we grouped brain samples by age. In each group, we randomly selected a *reference* right hemisphere. We registered the reference right hemispheres to each other in order of age, i.e. P1 → P0; P2 → transformed P1, etc. We further registered the remaining, non-reference right hemispheres to transformed reference hemispheres of the same age. We copied all transformations to the mirror reflected images of the left hemispheres. Thus, we put all hemispheres in the dataset to common coordinates.

Based on these registrations, we computed a distance matrix reflecting pairwise differences between *all* hemispheres in our dataset (correlation distances between *filtered* images; *Figure 5A*). We used this distance matrix in the CMDS dimensionality reduction algorithm (Torgerson, 1952) to compute the adjusted ages of each hemisphere in the dataset based on its developmental stage (*Figure 5B*). Specifically, for each hemisphere, we computed a coordinate along the 1^st^ CMDS dimension. We used these CMDS coordinates in linear regression to estimate the adjusted ages of hemispheres (*Figure 5B*). We checked whether the differences between the adjusted ages of two hemispheres within the same brain are small (*Figure 5D*).

### Display

To reconstruct the dynamics of brain development, we generated an average-case brain image for every time point within the span of the samples’ *adjusted ages* (*Figure 6D*) as follows. The weight (contribution) of a given brain hemisphere to a given time point was defined by a Gaussian curve with the maximum at the *adjusted age* of the hemisphere and the standard deviation equal to 1/2 day (1/2 of the age step between the groups).

At every time point, we normalized the sum of the weights of all hemispheres to one (Figure 6C). We used these normalized weights to combine (sum) all images of the hemispheres at every time point. As the normalized weights changed over time, the contributions of different brains to different time points varied forming a 3D animation of transitions between average-case developmental stages. Although our data were registered to a single hemisphere, we appended the animation with its mirror reflection to display the (symmetric) dynamics in the whole brain. To display the brain growth, we performed the linear fit of the brain sample sizes (Figure 6A, B) and scaled the averagecase images accordingly. To reconstruct the differences over time, for every time point we computed a differential image between the average-case images at the current time point and at the current time point minus one day. We color-coded the positive/negative changes in the intensity using red/blue color respectively (*Figure 6E*).

### Sample data

We illustrated our algorithm using 3D images of 28 whole mount perinatal mouse brain samples (P0 – P9) stained with 5-ethynyl-2’-deoxyuridine (EdU) to reveal dividing cells (Lazutkin et al., 2019) and imaged with a custom Olympus MVX10-based light-sheet microscope (Morozov et al., 2010) at the Z-resolution of 12 μm and the XY-resolution of 4 μm. The runtime of our algorithm was under 5 minutes per brain on a personal computer (Dell Precision 7510 using 7 GB of RAM and 1 CPU core at 3.3GHz). We verified the alignment by visually examining the comprehensive set of 2D slices for all aligned pairs of brain hemispheres (*Figure 3C,D*; Figure *4*).

**Figure 4.**
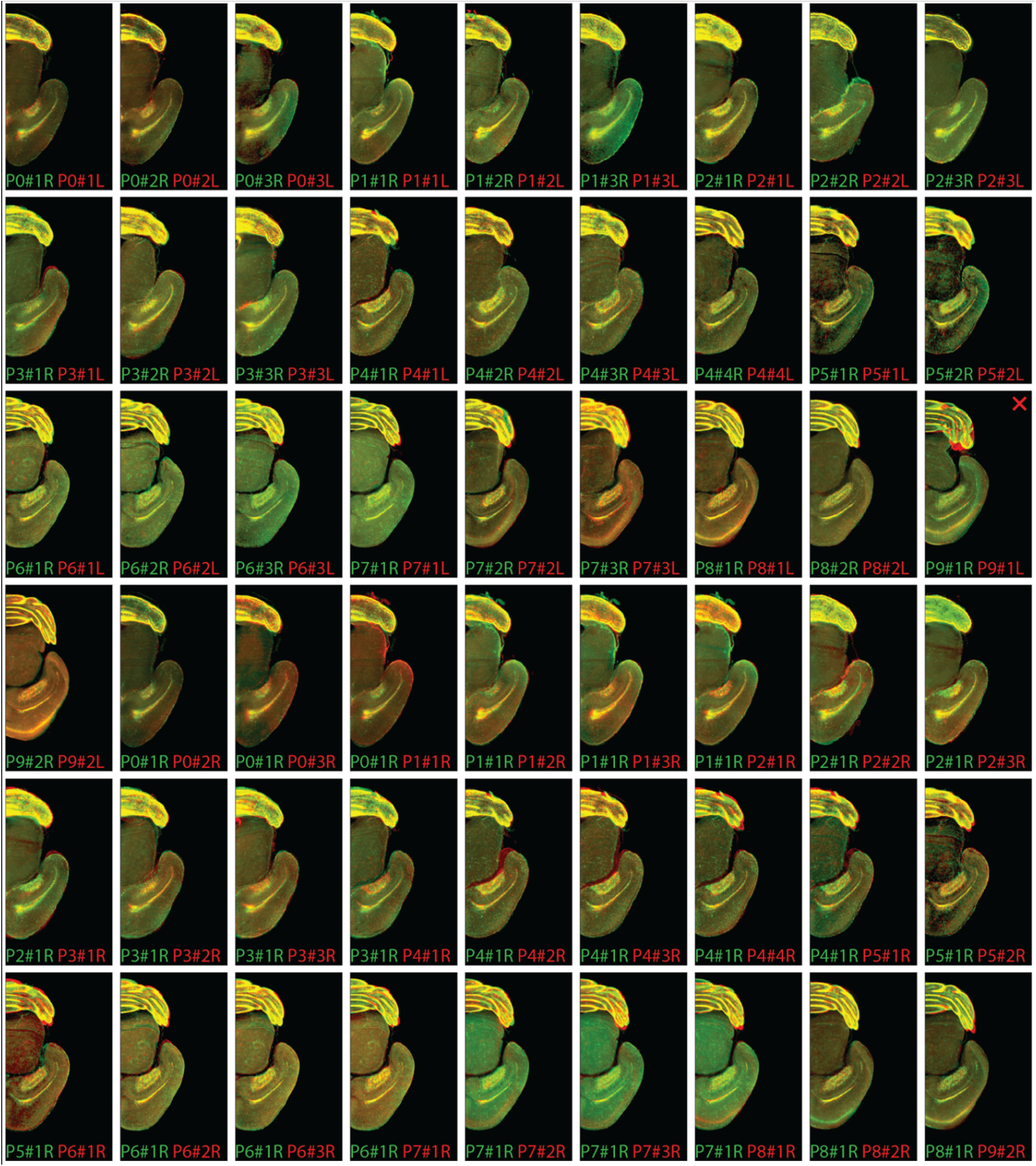
Registration was successful for 53 of 54 pairs of hemispheres from 28 whole-mount samples of developing brains in the testing set. Top: 28 pairs of left / right hemispheres from the same brains aligned to each other (27 successful). Bottom: 26 pairs of pre-registered hemispheres from 27 different brains (the 28^th^ brain with misaligned left /right hemispheres was not used here). For each aligned pair, we show a representative 2D optical section to illustrate the match of individual brain regions and highlight the potential differences. The 1 unsuccessfully registered pair of hemispheres is marked with the “X” sign (see cerebellum).

**Figure 5.**
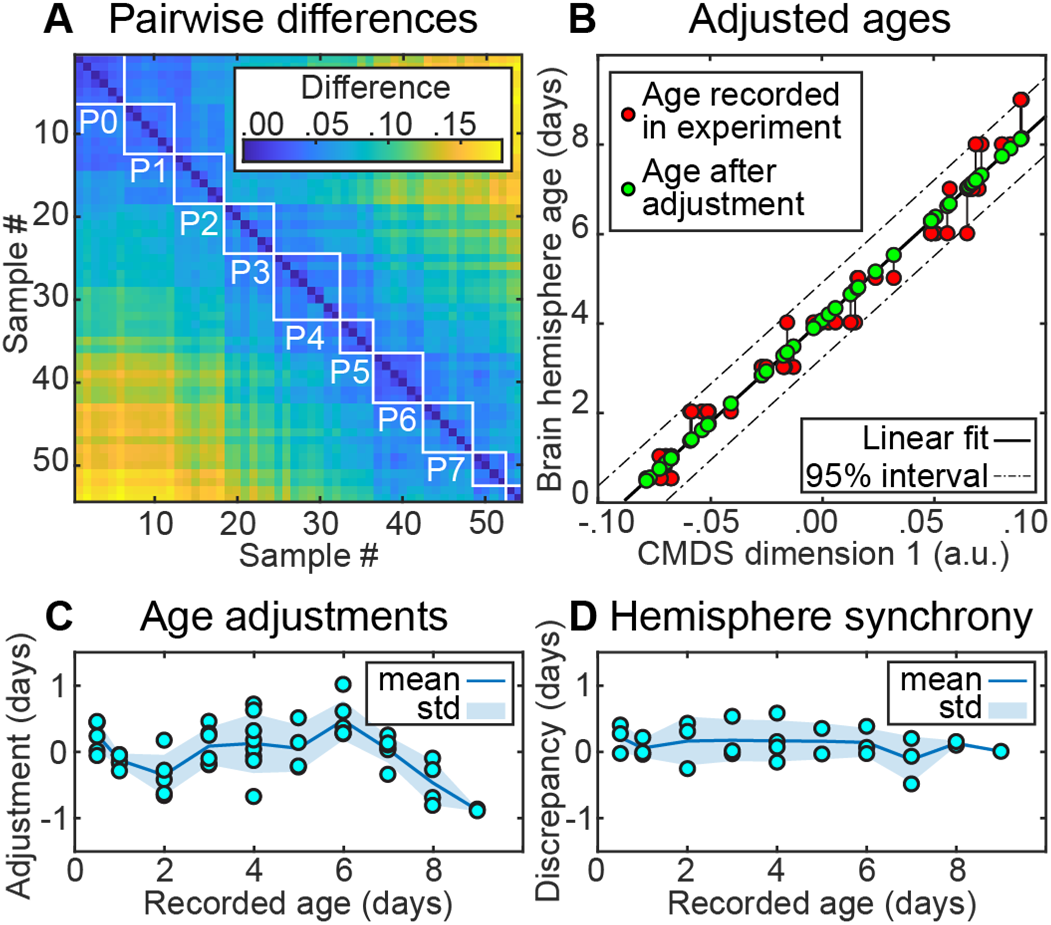
Temporal brain registration. We used differences between spatially registered brain samples to adjust the estimates of their developmental ages. (A) Pairwise differences (one minus Pearson correlation) between filtered 3D images of 54 hemispheres (of 27 brains). Boxes outline samples of the same ages as recorded in the experiment (P0; P1; P2 etc.). (B) Ages of brain samples as a function of the first CMDS component: the ages recorded in the experiment (red), linear fit (solid line) and the adjusted ages (green), 95% confidence interval (dashed line). (C) Age adjustment (difference between the adjusted and recorded ages) does not exceed one day. (D) Hemisphere synchrony (discrepancy between the adjusted ages of the left and right hemispheres in each brain).

**Figure 6.**
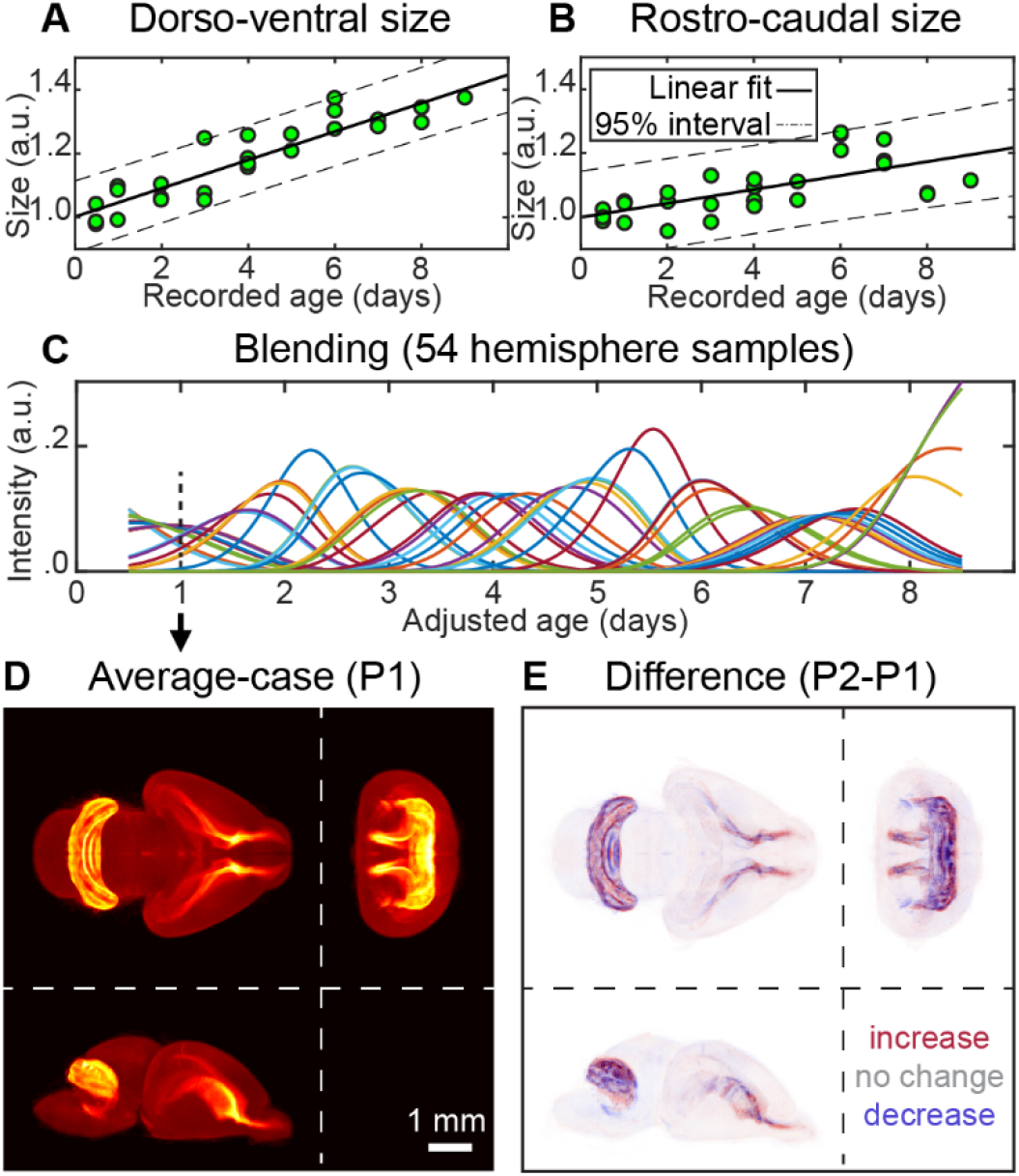
Using spatially and temporally registered brain samples to observe continuous brain development dynamics in 3D. (A) Dorsoventral and (B) rostrocaudal sizes of the brains normalized to the size at birth. Linear fit (solid line) and 95% confidence interval (dashed line). (C) Blending of 54 hemisphere samples. Colored lines show weighting contributions of each hemisphere to every time point. Contribution (or intensity) is maximal at the sample’s adjusted age; it decays with a standard deviation of 1/2 of the sampling rate (1/2 day); total intensity adds up to one at every time point. (D) Weighted average image for P1 developing mouse brain in accordance with weighting curves (C). (E) Difference between weighted average images of P2 and P1 mouse brains. The increases and decreases in cell proliferation (EdU+ cell density) are color-coded by the intensities of red and blue respectively.

## RESULTS

The focus of this work was on designing algorithms for reconstructing developmental dynamics of the perinatal mouse brain via the registration of brain samples in space and time. Several software packages are available for spatial registration of brain samples. NiftyReg package (Modat et al., 2010) developed for and used by the fMRI community and its neuroscience-oriented derivatives such as the aMAP (Niedworok et al., 2016) offer state-of-the-art registration for adult mouse brains. Registration of perinatal brains, however, brings new challenges. First, perinatal brains undergo significant transformations in a short time. They grow and change their overall shape as individual brain regions continue to develop. Thus, an algorithm for perinatal brain registration needs to overcome the pronounced shape differences between developing brains. Second, there is uncertainty in what a perinatal brain should look like at an arbitrary moment of development, because current brain atlases focus on major stages of development only (Chuang et al., 2011). Consequently, registration of perinatal brains must keep a balance between the precision of alignment and preservation of the original data. Third, the developmental unfolding of each brain proceeds at its own pace. As a result, algorithms for perinatal brain registration need to compensate for individual development paces by registering the brains in time. Below, we describe our methods to register diverse perinatal brains (Figure *1*). These methods aim to: i) maximize registration precision in context of dissimilar brain samples, ii) preserve the original data to the greatest extent possible, and iii) account for individual developmental paces. We illustrated each step with examples from 28 whole mount perinatal brain samples (P0 – P9) stained against dividing cells with 5-ethynyl-2’-deoxyuridine (EdU, (Lazutkin et al., 2019).

### Extracting brain region contours yields better registration

High precision of registration is important in studies involving perinatal brains. Brain samples representing distant developmental stages may be largely dissimilar, i.e. they have only a small number of common landmarks, resulting in unreliable registration. Conversely, brain samples representing adjacent developmental stages may be similar, and thus may be reliably registered in sequence. Because alignment errors accumulate over time, it is important to maintain high precision of registration. On the mesoscale level, a precise registration implies the perfect match of individual anatomical structures. In brain atlases, the anatomical structures are delineated using tissue borders (Chuang et al., 2011) in accordance with morphological staining (Paxinos and Franklin, 2004) or autofluorescence (Niedworok et al., 2016). Thus, the requirement for the precise mesoscopic-level registration can be reformulated as a necessity to match the *contours* of anatomical structures.

Aligning the contours of anatomical structures, besides being our immediate objective, also facilitates precise registration. For both raw images and structure contours, the similarity between two brain images is maximized when two brains are perfectly aligned. In the case of a slight displacement, raw images would still retain a massive overlap, keeping similarity almost unchanged. In contrast, displaced structure contours overlap little, making two images less correlated. We, therefore, expect contour-based registration to be more sensitive to structure mismatches when compared to conventional raw-imagebased approaches (Li et al., 1995).

The same logic, however, predicts difficulty in finding the initial coarse alignment in contour-based registration. Because the contours of unaligned brains are weakly correlated, the problem of aligning them becomes highly non-convex – and consequently much more difficult to solve. To facilitate the initial coarse alignment while retaining the benefits of a subsequent fine alignment, one may consider registration of the brain area contours *combined* with the binary mask of the sample (*Figure 2D*).

To test these arguments, we performed registration of a pair of different perinatal brains using raw images (*Figure 2E*), brain region contours (*Figure 2G*), and a combination of the brain region contours with the binarized images obtained after thresholding the raw images using 1% of maximum intensity as a threshold (*filtered* image, *Figure 2H*). To extract contours of morphologically defined structures in perinatal brains (Figure 2C), we used the Laplacian of Gaussian (LoG) filter (Huertas and Medioni, 1986). We also performed registration using the binarized images (*Figure 2F*) for completeness. As expected, the *raw-image-based registration* resulted in an adequate alignment, although some of the brain areas were misaligned (*Figure 2E*). The misalignment, despite being relatively small, was impactful in areas such as the olfactory bulb (OB). In the particular example of P5-P6 brains shown in *Figure 2E*, the image intensity was dominated by the cerebellum (see Figure *4*), and therefore the Ventricular-Subventricular Zone (V-SVZ) and the Rostral Migratory Stream (RMS) regions had a relatively small impact on the correlation between the source and the target brain images. Moreover, the OBs were mechanically flexible relative to the rest of the brain and, therefore, were distorted during chemical treatments of the samples. Mechanical flexibility paired with the low impact on the overall correlation between samples made the OBs an error-prone region in brain registration and, thus, required additional processing of raw images.

Based on these considerations, in this work, we used an image preprocessing step which combined extracted brain region contours with the binary masks of the samples. Registration performed on such preprocessed data resulted in better alignment of the 3D brain images (*Figure 2H*). Specifically, in two aligned images we observed significant overlap between the OBs and RMSs (*Figure 2H*). Both components of preprocessing were necessary: in separate *binary-mask* and *contour-based registrations* we observed large mismatches between resulting images (*Figure 2F,G*). Overall, a sum of the binarized image and the contours extracted using the LoG filter yielded the best alignment quality among the tested approaches.

### Attention-gated simulated annealing algorithm yields fast and robust registration

Algorithms for brain registration are expected to yield reliable alignment despite variability in samples. This requirement is especially important for perinatal (developing) brains. In the course of development, brain landmarks evolve: the brain grows and changes shape as brain regions continue to be fully formed. Below, we argue that simulated annealing (Van Laarhoven and Aarts, 1987), a Monte Carlo algorithm designed to find optima of non-convex functions, is well-suited to address such variability in samples. We illustrate the algorithm’s performance using 28 samples of perinatal mouse brains.

Pronounced differences in samples are challenging for automatic registration. Many optimization methods used for image registration (e.g. Powell’s, Gauss-Newton, Nelder– Mead, gradient descent, etc.) may require modifications in the conditions of non-convex measures of image similarity (Oliveira and Tavares, 2014). Such modifications include convexifying the task via registering images at different resolutions (Oliveira and Tavares, 2014) or overcoming local similarity minima by varying the optimizer’s step (Klein et al., 2009b). Such adjustments are necessary because large-scale displacements in brain samples may require correspondingly large steps of the optimizer, but at the same time, larger steps of the optimizer may result in the divergence of an algorithm. In simulated annealing, the problem of non-convex maximization is solved by allowing transient decreases in the objective function (proportional to similarity between the images; Figure 3E).

This way, the algorithm is equipped to escape from local maxima of cost function and to overcome pronounced differences between samples by taking multiple steps which can lead to both its decrease and increase (Van Laarhoven and Aarts, 1987). The algorithm is biased towards an overall increase in the cost function (similarity), which facilitates finding its regional maxima.

Alongside the benefits of simulated annealing, we should consider its potential disadvantages. Monte Carlo methods – the broader class of algorithms – are generally efficient when the number of variables to be optimized does not exceed 100 (Qian et al., 2016). In our setting, the algorithm needs to operate on a grid of up to ~400 nodes in 3D space. Nevertheless, this seemingly excessive number of nodes was not a concern in our approach for the following reasons. First, the registration is local, e.g. displacements of the transformation grid nodes in the cerebellum do not interact with the registration of the olfactory bulb. Second, we implemented the attention-gated alignment mechanism that selected the nodes to be altered in proportion with dissimilarity in their neighborhoods (see Methods). Finally, at every iteration, we only updated the content of image transformation grid cells adjacent to the displaced node. We, therefore, expected a reasonable convergence rate of simulated annealing in our brain registration task.

We first show that the ability to decrease the objective function is important for brain alignment. To do so, we compared our simulated annealing algorithm (Figure 3G-I), with the greedy algorithm produced by setting the temperature in the simulated annealing procedure to zero (Figure 3F). The greedy algorithm thus prohibited transient decreases in the objective function during optimization. For comparison, we used hemispheres of P5 and P6 brains, where the P5 brain was preliminarily registered to a P4 → … → P0 reference brain. This is an extreme example intended for illustration purposes; in our algorithm, we register brains in the order that makes differences between brains less pronounced (see below). We show that the greedy algorithm did not succeed in registering dissimilar brain samples (e.g. mismatch in the hippocampus in Figure 3F), whereas our attention-gated simulated annealing algorithm yielded sufficient overlap between fine brain structures (Figure 3I). We further show that our attention mechanism, choosing the grid nodes for adjustment based on dissimilarity in their neighboring cells of the grid, improved the algorithm’s convergence. In the example where all grid nodes were selected with equal probability (no attention; Figure 3H), the algorithm did not converge to satisfactory alignment given the same number of iterations. Finally, we show that discounting the objective function with deformation energy was also important for algorithm convergence. When undiscounted, simulated annealing deviated from optimal solution significantly, resulting in pronounced mismatches of the brain structures (e.g. hippocampus in Figure 3G). These observations suggest that using simulated annealing, attention-gating and discounting the objective function with deformation improves registration of 3D images of the brain.

We then tested the robustness and convergence rate of simulated annealing in the brain registration task. We performed registration of 28 whole mount perinatal mouse brain samples (P0 – P9), sampled daily, stained against dividing cells, and pre-filtered as described in the section above (Figure 2D). We split the brains into left and right hemispheres, registered the hemispheres from the same brain to each other, and registered pairs of right hemispheres from different brains. To save computing time and to eliminate the need to align dissimilar samples, we did not register all pairs of right hemispheres but only selected samples from the nearest time points (see Methods for rules of pair formation and registration order). We verified the alignment using the full collection of virtual slices. The match of fine brain structures – including cerebellum layers and the RMS – was observed in 27 out of 28 samples transformed to their reflections (*Figure 4*, top half) and then in 26 of 26 right hemispheres registered sequentially (*Figure 4*, bottom half). In each instance, registration took under 5 minutes per pair of brains on a laptop.

Overall, our results indicate that attention-gated simulated annealing performed on pre-filtered images is an efficient tool for spatial registration of brain samples. The algorithm is not computationally demanding and can be deployed on a personal computer. It is robust to variability in samples, including diverse sets of samples of perinatal brains.

### Temporal brain registration using the CMDS algorithm helps reduce variability in developmental dynamics

Perinatal brains of the same age display differences in shapes, sizes and developmental patterns, e.g. spatial distributions of dividing cells. Such variability may obscure the underlying developmental process and needs to be compensated for by an alignment algorithm. Below we propose a way to account for this variability.

To quantify the potential sources of variability in developing brains, we performed correlation analysis of the *filtered* brain images (mask + contours; Figure 2D) in 27 (out of 28) well-registered samples of perinatal mouse brains (P0 – P9) separated into 54 individual hemispheres. We noticed that some of the P1 brains looked like typical P0 brains; some of the P4 brains resembled P3, etc. (Figure 5A). Similarities between a fraction of the brains of different ages implied that some variability in the brains could be explained by temporal displacements in their development. We further reasoned that significant displacements were especially likely to be observed in studies of the perinatal brain where samples are dated with respect to birth – an event only approximately related to brain development. Thus, Chuang et al. (2011) used the date with respect to conception, not birth, to define brain development stages. Overall, if temporal shifts in development underlie anatomical variability, accounting for these shifts may unmask fine details of development otherwise averaged out.

To account for temporal shifts in brain development, we implemented *temporal registration* of brain samples. We used classical multidimensional scaling (CMDS), a linear dimensionality reduction technique (Torgerson, 1952), to “synchronize” brain samples by adjusting their ages. CMDS algorithm placed similar brains close to each other in time and placed the different ones apart based on the degree of their overall anatomical dissimilarity (Figure 5B). Unlike nonlinear embedding techniques such as Isomap (Tenenbaum et al., 2000), CMDS relied on both small and large-scale differences between samples. In our routine, relying only on small differences between samples for temporal registration could highlight artifacts induced by the order of spatial registration. This is because all brains within a same-age group were registered to one reference brain and imperfections of in-group registration were smaller than those across the groups. Overall, we expected temporal registration with CMDS algorithm to resolve uncertainty in brain development pace and to uncover refined dynamics of brain development.

To test the above arguments, we performed temporal registration of 27 perinatal mouse brains (54 hemispheres). First, we show that the age adjustments using CMDS did not exceed one day and did not increase over time (Figure 5C). The adjustments roughly corresponded to the uncertainty in the duration of mouse pregnancy (Dewar, 1968). At the same time, we observed no significant differences between the adjusted ages of left and right hemispheres within the same brains (Figure 5D). The 1^st^ CMDS dimension explained 94.5% of the variance in the embedding. These observations suggest that dating the samples relative to birth may be a major source of observed anatomical variability in development of perinatal brains.

Determination of the samples’ developmental age allowed us to monitor developmental dynamics in an ‘average’ brain. To this end, we first distributed the registered brains on the timeline in accordance with their adjusted ages (Figure 6C). Then we built the representation of the average brain by combining the aligned observed samples at each time point using a set of Gaussian weights. We then were able to both monitor the ongoing brain dynamics and to determine changes occurring in the distribution of EdU+ cells (Figure 6D,E; Figure *7*). As a result, day-to-day variability in the samples (L1 norm of the daily differences over the voxels in average 3D images downsampled to 1/8 of the original resolution) decreased by 31%. This suggests that temporal shifts in development are the important contributors to anatomical variability in developing brains defined on mesoscale and CMDS is efficient in estimating such shifts.

**Figure 7.**
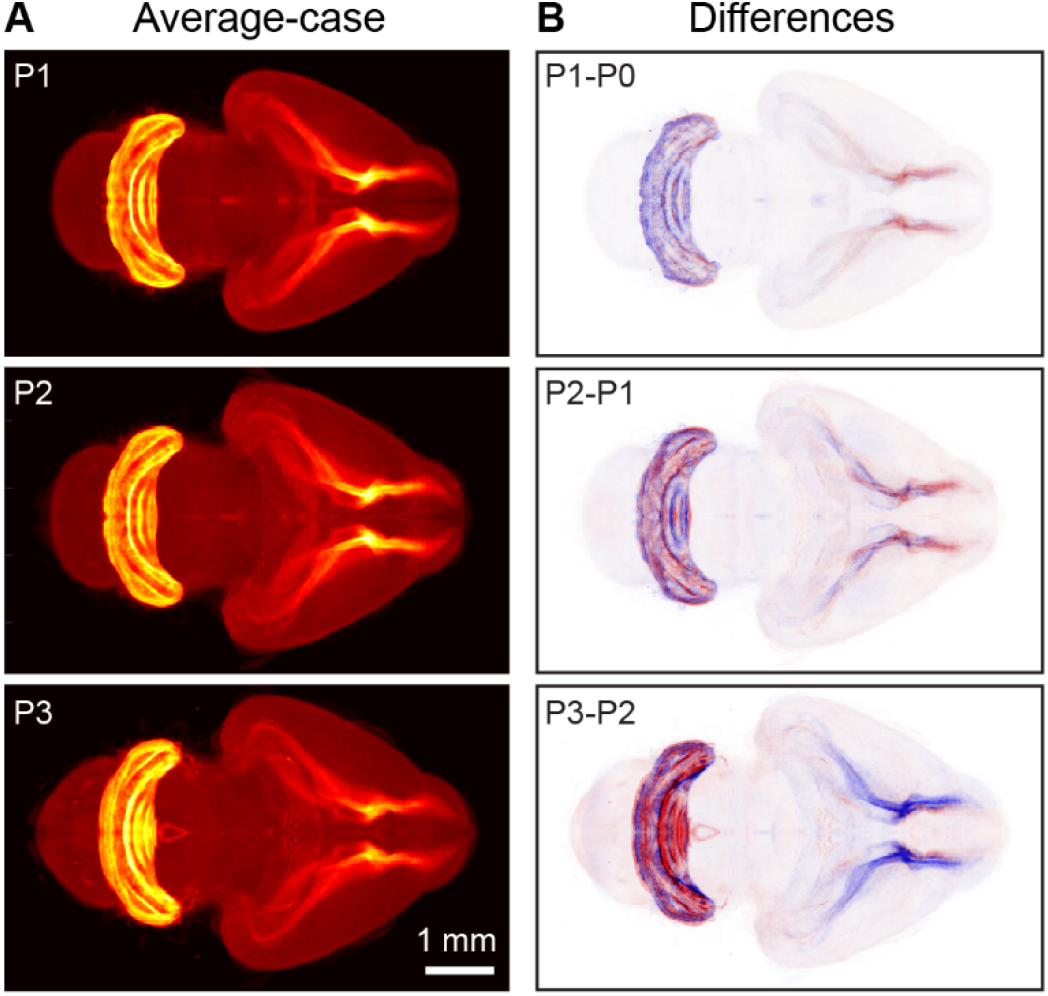
Example of continuous brain development dynamics detected using our algorithm. (A) Weighted average images for mouse developing brain on postnatal days P1-P3. (B) Average-case differences between P1-P3 and P0-P2 brains respectively reveal the dynamics of postnatal brain development. Over the course of these three days, the density of EdU+ cells in the cerebellum increases (blue to red) whereas it decreases in the RMS (red to blue). Differential images (B) highlight the development dynamics not easily noticeable in the weighted average images (A).

Overall, we conclude that the variability observed in perinatal brain samples may be partially explained by temporal shifts in brain development. Such shifts may arise due to a discrepancy between the moment of birth and the developmental stage of the brain at the time of analysis. These shifts can be estimated using the CMDS algorithm and – once accounted for – may reduce variability in the dynamics of brain development. We expect that data corrected this way may allow uncovering additional details of brain development dynamics.

## DISCUSSION

In this work, we have proposed a computational pipeline for reconstructing mesoscale dynamics of the developing mouse brain. We focused on the types of data which can only be collected *ex vivo*; therefore, we used multiple brains to infer the development dynamics. To combine the data, we proposed aligning 3D images of different brains (image registration). For reliable reconstruction of the development dynamics, we required high-precision alignment of variable brain samples. First, we showed that high-precision alignment can be achieved by using the contours of brain regions (Figure 2) instead of the raw images. We then showed that contours can be efficiently aligned using simulated annealing (Figure 3). This way, we combined the accuracy of feature-based registration approaches with the throughput of free-form approaches. We then used 28 samples of perinatal mouse brains at different developmental stages to show that our registration algorithm is robust to variability in samples (Figure *4*). Finally, we showed that individual paces of brain development can be accounted for by additionally registering brain samples in time (Figure 5), thus smoothing (denoising) transitions between developmental stages (Figure 6). Overall, the steps above enabled us to uncover developmental dynamics in perinatal mouse brains by using static images at different developmental stages (Figure *7*).

Reconstructing developmental dynamics from series of *ex-vivo* samples has several advantages compared to *in-vivo* imaging. First, *ex vivo* studies allow combining substantial imaging volume with high resolution. The best alternative, functional ultrasound imaging, allows to image the entire mouse brain *in-vivo* at the resolution of 100 μm (Macé et al., 2011). Alternatively, three-photon microscopy enables *in vivo* imaging at a cellular resolution up to the depth of 1300 μm (Horton et al., 2013). For multi-day imaging typical for developmental studies, both functional ultrasound imaging and three-photon imaging may require image alignment. At the same time, *ex vivo* brain samples allow obtaining cellular resolution in the entire brain (Chuang et al., 2011). Using *ex vivo* imaging also enables the broader choice of reporter molecules, such as various fluorescent labels (Susaki and Ueda, 2016).

Our pipeline is robust to potential inaccuracies of the individual algorithms used. Although each subroutine of the pipeline improved registration quality, together, these algorithms play redundant roles. For example, if raw images are used for registration instead of the filtered ones, the low-variance regions such as V-SVZ-RMS may still be aligned well because of the attention mechanism in simulated annealing. Should the brain area contours be highlighted too much compared to the background, the algorithm may not get trapped in an erroneous local maximum of similarity because of temperature in simulated annealing – which allows a transient decrease in the similarity between samples. Overall, we argue that the steps of our algorithm, when combined, lead to robust registration of brain samples.

Conventionally, the scope of registration was limited to pairs of samples (Prince et al., 2015). In this work, we have proposed a procedure for multi-sample registration, in both space and time. Our procedure allows one to use separate brain samples to reconstruct continuous dynamics of developmental processes and to trace related long-term changes (Figure *7*). The ability to capture developmental dynamics based on static data snapshots is especially important when the data can be only collected *ex vivo*. At the same time, our procedure supports all conventional use cases for registration algorithms, including the direct comparison of individual samples/groups and registration to common coordinate frameworks, e.g. (Allen Institute, 2017). In particular, our procedures can be combined with cell detection software (e.g. Renier et al., 2016; Shuvaev et al., 2017). For conventional applications, our procedures offer high registration quality and fast convergence rates. Finally, the procedures described in this paper are modular. Depending on the task, its parts (feature extraction, spatial registration, temporal registration, data display) can be used together, separately, or in combination with other packages of the user’s choice. The described algorithms can be downloaded at http://github.com/koulakovlab/registration.

## ACKNOWLEDGEMENTS

*This study was supported by grants from the National Institute of Aging (R01 AG057705 and R21 AG063004) and the National Institute of Mental Health (R21 MH118991) to G.E., grant from the National Institute on Drug Abuse (R01 DA050374) to A.K., grants from the Russian Science Foundation (17-15-01426 and 19-15-00247), Russian Foundation for Basic Research (19-29-04173) Ministry of Science and Higher Education of the Russian Federation (075-15-2020-801), and the Swartz Foundation. All manipulations with animals were carried out in compliance with the national and international Guides for the Care and Use of Laboratory Animals*.

